# Dram1 confers resistance to *Salmonella* infection

**DOI:** 10.1101/2021.03.21.436194

**Authors:** Samrah Masud, Rui Zhang, Tomasz K. Prajsnar, Annemarie H. Meijer

## Abstract

Dram1 is a stress and infection inducible autophagy modulator that functions downstream of transcription factors p53 and NFκB. Using a zebrafish embryo infection model, we have previously shown that Dram1 provides protection against the intracellular pathogen *Mycobacterium marinum* by promoting the p62-dependent xenophagy of bacteria that have escaped into the cytosol. However, the possible interplay between Dram1 and other anti-bacterial autophagic mechanisms remains unknown. Recently, LC3-associated phagocytosis (LAP) has emerged as an important host defense mechanism that requires components of the autophagy machinery and targets bacteria directly in phagosomes. Our previous work established LAP as the main autophagic mechanism by which macrophages restrict growth of *Salmonella* Typhimurium in a systemically infected zebrafish host. We therefore employed this infection model to investigate the possible role of Dram1 in LAP. Morpholino knockdown or CRISPR/Cas9-mediated mutation of Dram1 led to reduced host survival and increased bacterial burden during *S*. Typhimurium infections. In contrast, overexpression of *dram1* by mRNA injection curtailed *Salmonella* replication and reduced mortality of the infected host. During the early response to infection, GFP-Lc3 levels in transgenic zebrafish larvae correlated with the *dram1* expression level, showing over two-fold reduction of GFP-Lc3-*Salmonella* association in *dram1* knockdown or mutant embryos and an approximately 30% increase by *dram1* overexpression. Since LAP is known to require the activity of the phagosomal NADPH oxidase, we used a *Salmonella* biosensor strain to detect bacterial exposure to reactive oxygen species (ROS) and found that the ROS response was largely abolished in the absence of *dram1*. Together, these results demonstrate the host protective role of Dram1 during *S*. Typhimurium infection and suggest a functional link between Dram1 and the induction of LAP.

## Introduction

Autophagy has long been known as a fundamental housekeeping process wherein dysfunctional cellular components are captured inside double membrane vesicles that fuse with lysosomes to degrade and recycle the contents (Levine and Klionsky, 2004; Mizushima, 2007). More recently, autophagy has also become recognized as an integral part of the immune system, functioning not only as a direct anti-microbial mechanism but also contributing to regulation of the immune response (Deretic et al., 2013). Autophagy can function as a non-specific bulk process or as a selective mechanism mediated by receptors that recognize molecular degradation signals like ubiquitin (Boyle and Randow, 2013). The selective autophagy of invading microbes, referred to as xenophagy, is an innate immune effector mechanism targeting invading microbes either when present inside membrane-bound compartments or when they escape into the host cytosol (Huang and Brumell, 2014). A hallmark of all forms of autophagy is the conjugation of microtubule-associated protein 1 light chain 3 (MAP1LC3, hereafter LC3) with phosphatidylethanolamine on the autophagosomal double membrane structures (Mizushima et al., 2004). However, the autophagy machinery can also recruit LC3 to single membrane compartments, specifically phagosomes, in a process known as LC3-associated phagocytosis (LAP) (Sanjuan *et al*., 2007; Martinez, 2018; Upadhyay and Philips, 2019). Both xenophagy and LAP have been shown to be important for resistance to infections, but pathogens have also evolved mechanisms to subvert these host autophagic defenses (Huang and Brumell, 2014; Mostowy, 2013; Deretic *et al*., 2006; Hubber *et al*., 2017; Prajsnar *et al*., 2020).

There are various mechanisms by which autophagy can be induced in response to internal and external stress factors, such as nutrient restriction, DNA damage, and microbial invaders. One factor implicated in the activation of autophagy is DNA damage regulated autophagy modulator gene 1 (*DRAM1*), which encodes a member of an evolutionary conserved family of six transmembrane proteins (Mah et al., 2012). *DRAM1* was first reported as direct target gene of the tumor suppressor protein p53 and shown to play a role in p53-mediated autophagy and apoptosis (Crighton *et al*., 2006). DRAM1 has been implicated in several types of cancers (Galavotti *et al*., 2013; Chen *et al*., 2018; Meng *et al*., 2018). Overexpression of *DRAM1* has been shown to increase basal levels of autophagosome numbers, indicating that DRAM1 can act early in the autophagy process, contributing to autophagosome formation (Mah et al., 2012). However, the DRAM1 protein predominantly localizes to lysosomes and there is evidence also for functions at later steps in the process, showing that DRAM1 enhances autophagic flux and promotes ATPase activity and lysosomal acidification (Zhang *et al*., 2013; Geng *et al*., 2020). DRAM1 is thought to mediate crosstalk between autophagy and apoptosis by interacting with the proapoptotic protein BAX (Guan *et al*., 2015). Furthermore, in an autophagy-independent manner, DRAM1 interacts with transporter proteins to promote lysosomal amino acid efflux, which activates the nutrient-sensing complex mTORC1 (Beaumatin *et al*., 2019). Other molecular interactions of DRAM1 that could explain its mode of action at early as well as late steps of the autophagy pathway remain to be elucidated.

In addition to DNA damage, *DRAM1* is also induced by infection (Laforge *et al*., 2013; van der Vaart et al., 2014). In HIV-infected T-cells, DRAM1 has been shown to function downstream of p53, triggering lysosomal membrane permeabilization and cell death (Laforge *et al*., 2013). In contrast, we have shown that the induction of *DRAM1* in macrophages infected with *Mycobacterium tuberculosis* is independent of p53 and mediated instead by transcription factor NFκB, which functions downstream of pathogen recognition by Toll-like receptor (TLR) signaling (van der Vaart et al., 2014). Similarly, induction of the zebrafish homolog of *DRAM1* (*dram1*) by *Mycobacterium marinum* infection relies on the TLR adaptor molecule MyD88 (van der Vaart et al., 2014). *M. marinum*-infected zebrafish embryos develop a tuberculosis-like disease and we have shown that Dram1 plays a host protective role in this model (van der Vaart *et al*., 2014; Meijer and van der Vaart, 2014; Zhang *et al*., 2020). Specifically, we found that knockdown of *dram1* strongly reduces the co-localization of Lc3 with *M. marinum*, whereas overexpression increases Lc3-*M. marinum* co-localization in a manner dependent on the selective autophagy receptor p62 (Meijer and van der Vaart, 2014). Additionally, we observed that in *dram1*-/-CRISPR mutant zebrafish, not only Lc3 co-localization but also the acidification of *M. marinum-*containing vesicles was reduced (Zhang *et al*., 2020). Failing to contain the intracellular infection, macrophages in *dram1-/-*zebrafish succumbed to pyroptotic cell death (Zhang *et al*., 2020).

In the present study we sought to investigate the role of Dram1 during infection of zebrafish embryos with another intracellular pathogen, *Salmonella enterica* serovar Typhimurium (S. Typhimurium). A zebrafish model for *S*. Typhimurium infection has previously been established and *dram1* expression is inducible following intravenous injection of the pathogen in this model (van der Sar *et al*., 2003; Stockhammer *et al*., 2010). In our recent work, we have shown that LAP is the main autophagic process targeting *S*. Typhimurium following phagocytosis of the pathogen by macrophages in the zebrafish host (Masud *et al*., 2019a). Furthermore, we found that inhibition of LAP, by knockdown of factors specific for this process (Rubicon and NADPH oxidase), impaired host resistance (Masud *et al*., 2019a; Masud *et al*., 2019b). Here, we demonstrate that Dram1 is required for host resistance to *S*. Typhimurium infection and promotes Lc3 association with this pathogen, supporting a role for Dram1 in LAP-mediated host defense.

## Results

### Knockdown and overexpression of *dram1* indicates its function in host defense against *Salmonella* infection

In order to investigate the role of Dram1 during *Salmonella* infections we modulated levels of *dram1* by splice blocking morpholino knockdown or mRNA injection at the 1-2-cell stage. We challenged the host at 2 days post fertilization (2 dpf) with *S*. Typhimurium by intravenous injection and recorded progression of the resulting infection by survival curves and CFU determination **(Figure 1A)**. We found that *dram1* knockdown resulted in hypersusceptibility to *Salmonella* infection, where at 48 hours post infection (hpi) nearly 74 % of hosts succumbed to *S*. Typhimurium infection in significant contrast to 54 % of mortality in control hosts **(Figure 1B)**. Moreover, only 4 % of *dram1*-deprived hosts were alive at 72 hpi compared to 29 % of the control group **(Figure 1B)**. Additionally, under knockdown conditions of *dram1*, infected hosts contained significantly higher *S*. Typhimurium bacterial counts at 24 hpi **(Figure 1C)**. Conversely, the overexpression of *dram1* by mRNA injection resulted in higher survival rates **(Figure 1B)** and lower numbers of *S*. Typhimurium bacteria at 24 hpi **(Figure 1C)**. These results indicate that Dram1 restricts *Salmonella* infection.

**Figure 1:**
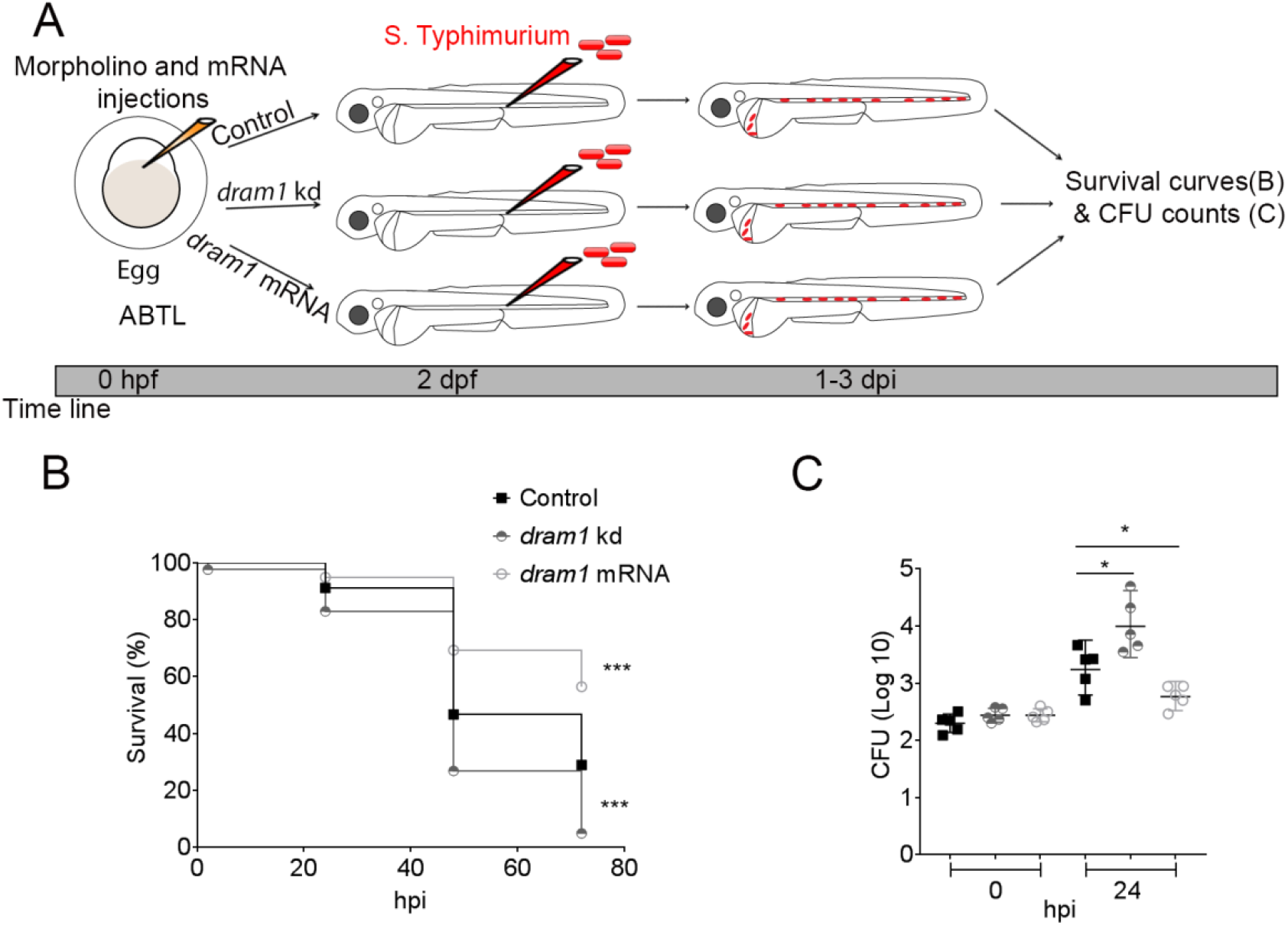
Dram1 is required for effective host defense during *Salmonella* infections. **A**: Workflow of experiments followed in B&C along with the time line of developing embryos. **B**: Survival curves of embryos with *S*. Typhimurium infection expressing different levels of *dram1*. Survival curves of control embryos were compared to *dram1* knockdown and overexpressing groups. One representative of three replicates is shown (n=50).**C:** CFU counts for infected larvae groups in B. For each of the three groups five embryos/larvae per time point were used and the log transformed CFU data are shown with the geometric mea mean per time point. One representative of three replicates is shown. Error bars represent SD. ***P< 0.001, *P< 0.05.

### Dram1 is required for the Lc3 response of the host that targets *Salmonella* phagocytosed by macrophages

We have previously shown that the autophagy response of macrophages towards *Salmonella* in our model occurs mainly as Lc3-associated phagocytosis (LAP) (Masud *et al*., 2019a). To study whether Dram1 contributes to the Lc3 targeting of *Salmonella*, we injected *dram1* morpholino into embryos of the *Tg(CMV:GFP-maplc3b1)* line (hereafter referred to as GFP-Lc3) and infected these with *mCherry*-expressing *S*. Typhimurium **(Figure 2A)**. We observed that GFP-Lc3 associations with *Salmonella* cells were significantly Dram1 dependent, since morpholino knockdown resulted in more than two-fold reduction of GFP-Lc3-positive infected phagocytes **(Figure 2B,C,E)**. Thus, the increased susceptibility of *dram1* knockdown embryos **(Figure 1)** is associated with diminished levels of Lc3-*Salmonella* associations. Next, we investigated if the positive effect of mRNA*-*mediated *dram1* overexpression on restricting infection **(Figure 1)** is accompanied by an increased Lc3 response towards the pathogen. Indeed, infection of control and *dram1* overexpressing GFP-Lc3 embryos showed a significant increase of GFP-Lc3 associations with *S*. Typhimurium bacterial cells in the overexpression group **(Figure 2 D and E)**. These results lead us to propose that Dram1 promotes the LAP response during *Salmonella* infections, resulting in a more effective host defense.

**Figure 2:**
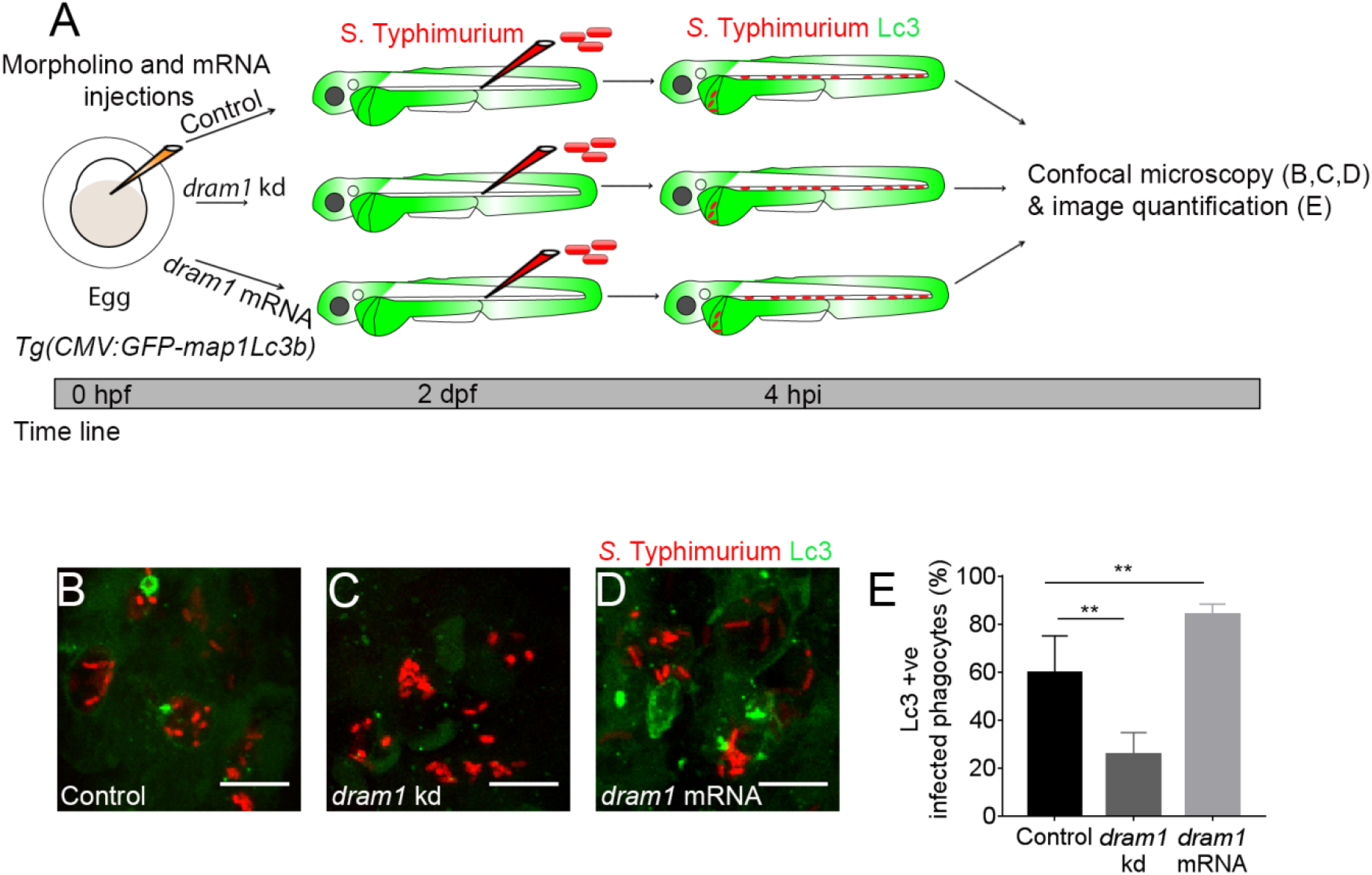
Dram1 modulates GFP-Lc3 associations with *S*. Typhimurium. **A:** Workflow and time line of experiments in B-E. **B-D:**. Representative confocal micrographs of *Tg(CMV:GFP-Lc3map1b)* embryos from control (B), *dram1* kd (C), and *dram1* mRNA (D) groups infected with *mCherry*-expressing *S*. Typhimurium at 4 hpi. **E:** Quantification of GFP-Lc3-*Salmonella* associations at 4 hpi. Five embryos per group were imaged over the yolk sac circulation valley and quantified as percentages of infected phagocytes positive for GFP-Lc3-*Salmonella* associations (LC3+ve) over the total number of infected phagocytes. Error bars represent SD. One representative of three replicates is shown. Scale bar (B-D) = 10μm. **P<0.01.

### Dram1 mutation recapitulates the infection phenotype of *dram1* morpholino-induced knockdown

In order to verify the results obtained using morpholino-mediated knockdown, we studied *S*. Typhimurium infection in a recently established *dram1* mutant line (Zhang *et al*., 2020). We infected *dram1* -/-and *dram1* +/+ embryos with *S*. Typhimurium **(Figure 3A)** and observed that *dram1*-/-hosts showed increased mortality during *S*. Typhimurium infection as compared to *dram1*+/+ controls **(Figure 3B)**. Bacterial growth determination confirmed the *Salmonella*-restricting role of Dram1 as significantly higher bacterial counts were retrieved from *dram1-/-*embryos at 24 hpi as compared to *dram1+/+* embryos **(Figure 3C)**. The *dram1* mutants were subsequently tested for Lc3 targeting of *Salmonella*. We observed significantly less GFP-Lc3 associations with *Salmonella* cells in *dram1-/-*embryos compared to *dram1+/+* embryos **(Figure 3D-F)**, further confirming our initial observations using morpholino knockdown. The consistent results obtained in Dram1-deficient hosts achieved by knockdown or stable mutation support the function of Dram1 in LAP-mediated host defense during *Salmonella* infections.

**Figure 3:**
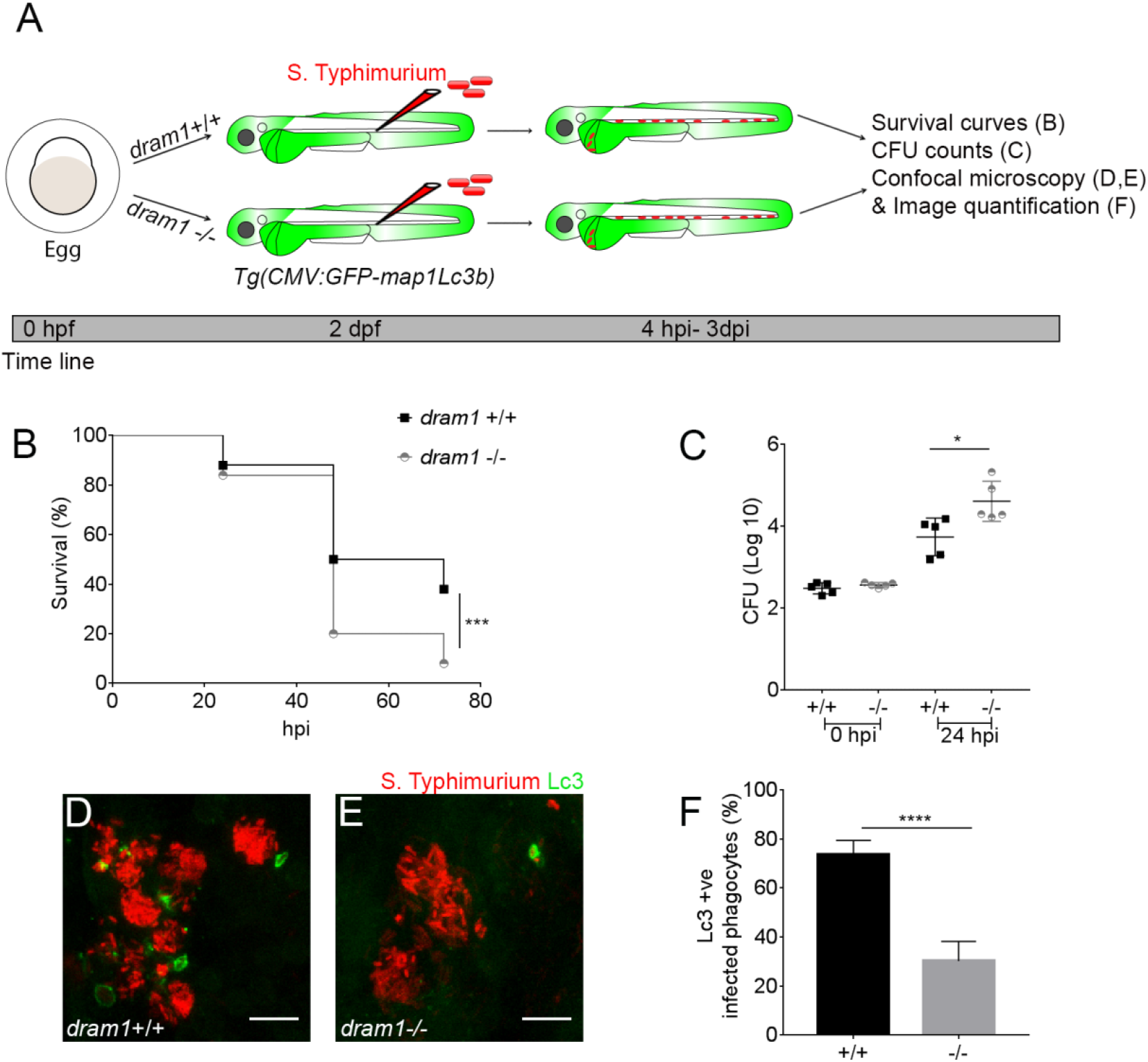
LAP induction in impaired in *dram1* mutants. **A:** Workflow and time line of experiments in B-F. **B:** Survival curves of *dram1*+/+ and *dram1*-/-embryos infected with *S*. Typhimurium. One representative of two replicates is shown (n=50). **C:** CFU counts recovered from *dram1* +/+ and *dram1* -/-infected individuals. Five embryos/larvae per time point were used and the log transformed CFU data are shown with the geometric mean per time point. One representative of three replicates is shown. Error bars represent SD. **D**,**E:** Representative confocal micrographs of GFP-Lc3-*Salmonella* associations in *dram1*+/+ (D) and *dram1*-/-(E) embryos in *Tg(CMV:GFP-Lc3map1b* background. **F**: Quantification of GFP-Lc3-*Salmonella* associations in *dram1*+/+ and *dram1* -/-embryos. Five embryos were imaged over the yolk sac circulation valley and infected phagocytes positive for GFP-Lc3-*Salmonella* associations were quantified as percentage over the total number of infected phagocytes. Error bars represent SD. One representative of two replicates is shown. Scale bar (D,E) = 10 μm, ****P<0.0001. **P<0.01, *P<0.05

### Dram1 is required for the phagocyte ROS response associated with LAP

LAP strictly depends on NADPH oxidase and Rubicon-mediated generation of ROS (Martinez, 2018). Therefore, to further study the link between Dram1 and the LAP response to *S*. Typhimurium, we determined the effect of Dram1 deficiency on reactive oxygen species (ROS) generation. To this end we utilized an *S*. Typhimurium ROS biosensor strain that contains a constitutively expressed *mCherry* reporter and a GFP reporter that is activated when bacterial cells are exposed to ROS (Burton *et al*., 2014) **(Figure 4A)**. We have previously shown that the activation of this ROS biosensor is strictly dependent on the expression of Rubicon and the NADPH oxidase component Cyba, demonstrating that this biosensor is a reliable indicator of the occurrence of LAP (Masud *et al*., 2019a). We observed that *dram1*-/-hosts were unable to activate the ROS biosensor in contrast to *dram1*+/+ individuals **(Figure 4B-D)**. Together, our results demonstrate that Dram1 deficiency impairs both GFP-Lc3 recruitment and ROS generation in infected phagocytes of the zebrafish host, consistent with the proposed role of Dram1 in the LAP response to *Salmonella*.

**Figure 4:**
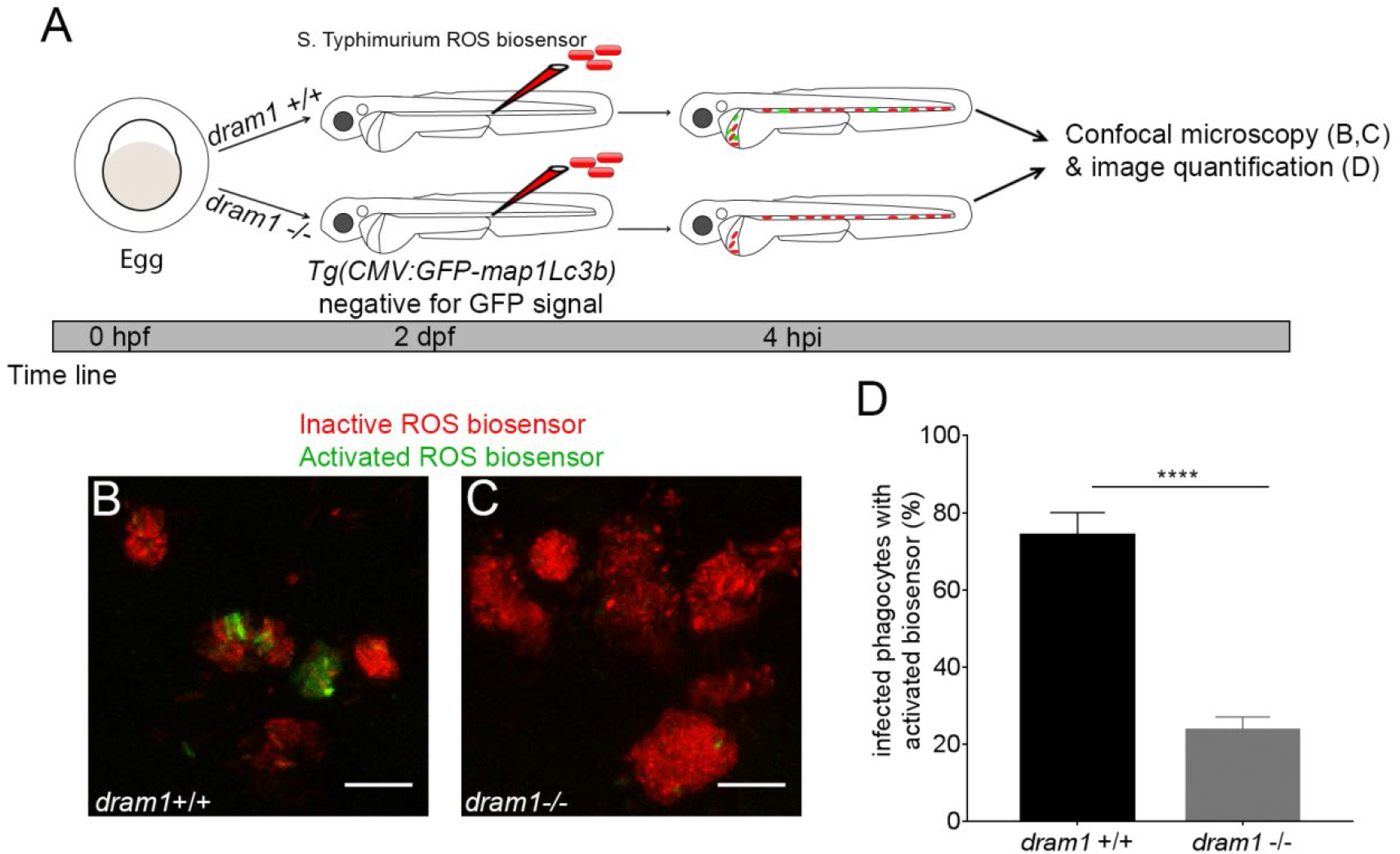
ROS generation is mediated by Dram1 during *Salmonella* infection. **A:** Workflow an time line of experiments in B-D. **B-C:**. Representative micrograph for *dram1*+/+ (B) and *dram1*-/-(C) where embryos were infected with a *Salmonella* biosensor strain for ROS. Inactive biosensor constitutively expresses mCherry and activated biosensor expresses GFP in addition to mCherry. Images were taken at 4 hpi. **D:** Quantification of ROS biosensor activation in *dram1*+/+ and *dram1*-/-at 4hpi. Numbers of phagocytes showing ROS biosensor activation (GFP and *mCherry* bacterial signals) or without ROS biosensor activation (*mCherry* bacterial signal only) were counted from confocal images and the percentages of ROS biosensor-positive over the total were averaged from five embryos per group. Error bars represent the SD. Scale bar (B-C) = 10 μm. ****P<0.0001.

## Discussion

In this study, we have used a zebrafish infection model to demonstrate that the autophagy modulator Dram1 promotes host resistance to systemic *Salmonella* infection. This work extends our previous finding that Dram1 mediates autophagic host defense in the zebrafish tuberculosis model. Importantly, our study implicates Dram1 in an infection context where LAP is the predominant autophagic host defense response.

LAP has recently been shown to function as a crucial host defense mechanism in bacterial as well as fungal infection (Martinez *et al*., 2015; Hubber *et al*., 2017; Sprenkeler et al., 2016; Masud *et al*., 2019a). In LAP, the autophagy marker LC3 is recruited to phagosomes by a mechanism that requires essential components of the autophagy machinery, but is independent of the ULK1 preinitiation complex (Martinez *et al*., 2015). DRAM1 has been shown to promote autophagy and autophagic flux, but has not previously been implicated in LAP (Crighton *et al*., 2006; Zhang *et al*., 2013). We have previously shown that macrophages in systemically infected zebrafish embryos target *S*. Typhimurium almost exclusively by LAP (Masud *et al*., 2019a). Lc3 recruitment to *S*. Typhimurium in zebrafish macrophages is dependent on two proteins that are essential for LAP: Rubicon and p22Phox, a component of the NADPH oxidase (Martinez *et al*., 2015). Depletion of either of these proteins abolishes GFP-Lc3 association with *S*. Typhimurium and prevents activation of a bacterial ROS biosensor gene (Masud *et al*., 2019a). Here, we show that both these responses are also inhibited by knockdown or mutation of *dram1*, suggesting that Dram1 is required for the targeting of *S*. Typhimurium by LAP. This Dram1-mediated process is host-protective, since we found that *dram1*-deficient zebrafish embryos are more susceptible to *S*. Typhimurium infection and that *dram1* overexpression promotes host resistance.

The molecular mechanism by which Dram1 promotes autophagic processes is currently unknown. Our data show that Dram1 is required not only for Lc3 recruitment but also for the ROS response in phagosomes. This suggests that Dram1 functions upstream of Rubicon, which has been shown to stabilize the NADPH oxidase. An attractive hypothesis is that Dram1 interacts directly with components of the Beclin1-VPS34-UVRAG-Rubicon complex to promote the association of Rubicon with phagosomes. This hypothesis is consistent with a recent study on a member of the human DRAM family, DRAM2, which has been shown to stimulate autophagy by disrupting the association of Rubicon with the Beclin1/UVRAG complex (Kim et al., 2011). The displacement of Rubicon relieves its inhibitory function from this complex and thereby promotes the activity of the class III phosphatidylinositol 3-kinase, VPS34, which is required for autophagosome formation and maturation. If Dram1 functions by a similar mechanism in the response to *Salmonella* infection, disruption of the Rubicon-Beclin1/Uvrag interaction would liberate Rubicon to associate with the *Salmonella*-containing phagosomes. This hypothesis would explain how Dram1 is able to stimulate autophagy and LAP at the same time. Another attractive hypothesis is that Dram1 stimulates the maturation of autophagosomes and LAPosomes by associating with lysosomal proteins, such as LAMP1 and LAMP2, similar to DRAM2 (Kim et al., 2011). Therefore, future work should focus on the effect of Dram1 on acidification of Salmonella-containing vesicles.

In addition to promoting LAP, it is likely that the activity of Dram1 can enhance other autophagic mechanisms against intracellular *Salmonella*, including the selective autophagy process that is mediated by ubiquitin receptors. Selective autophagy has been shown to function as an important defense mechanism against *Salmonella* in epithelial cells, which this pathogen enters by active invasion (Huang *et al*., 2009). Selective autophagy specifically targets cytosolic bacteria or detects membrane damage on the bacterial replication niche. In zebrafish, we have shown that this response is elicited by the escape of *M. marinum* from phagosomes and in this model Dram1 depends on the ubiquitin receptor p62 for its autophagy enhancing effect (van der Vaart et al., 2014). The above proposed interaction of Dram1 with Rubicon is consistent with positive effects on selective autophagy as well as LAP. In addition, the microbicidal capacity of Lc3-marked bacterial compartments might be enhanced by Dram1 activity, regardless of whether these compartments are of autophagosomal or phagosomal origin. This could be facilitated by delivery of ubiquitinated antimicrobial peptides through fusion with autophagosomes, which are present in increased numbers when Dram1 is overexpressed (Ponpuak *et al*., 2010; van der Vaart et al., 2014).

Different members of the DRAM family might have partially overlapping functions in host defense. In addition to a single copy of *dram1*, the zebrafish has two copies of the gene for Dram2, *dram2a* and *dram2b*. Based on our RNA sequencing data of leukocyte populations sorted from zebrafish larvae, *dram1* is the most abundantly expressed gene in macrophages, while the expression level of *dram2b* is at least 10-fold lower and *dram2a* is not detectably expressed (Rougeot *et al*., 2019). Furthermore, we have only found expression of *dram1* to be inducible by infection (Stockhammer *et al*., 2010; Benard et al., 2014). Our study did not provide any indication that other members of the zebrafish Dram family might be involved in LAP, since mutation of *dram1* abolished ROS biosensor activation in the *Salmonella*-containing phagosomes almost completely. However, the possible role of Dram2a/b in zebrafish host defense requires further investigation in the light of the recent report that *Mycobacterium tuberculosis* evades autophagy by inducing the expression of a microRNA (miR144*) that targets human *DRAM2* (Kim et al., 2017). This study has revealed that DRAM2 enhances antimicrobial activity in human monocyte derived macrophages, similar to the function that we have shown for Dram1 in the zebrafish host during infections with *M. marinum* and *S*.Typhimurium (van der Vaart et al., 2014); and this study). Therefore, accumulating evidence positions the members of the DRAM family as key players in autophagic defense against intracellular pathogens and it will be of great interest to further explore how these autophagy modulators might work in concert and how they could be therapeutically targeted.

## Materials and methods

### Zebrafish lines and maintenance

Zebrafish were handled in compliance with local animal welfare regulations and international guidelines specified by the EU Animal Protective Directive 2010/63/EU and maintained according to standard protocols (zfin.org). All studies were performed on embryos/larvae before the free feeding stage. Fish lines used for this study are AB/TL (wild type strain), transgenic lines *Tg(CMV:GFP-map1lc3b)* (He & Klionsky, 2010) *Tg(mpeg1::mcherry-F)*^*umsF001*^ (Bernut *et al*., 2014) and *dram1-/-*and *dram1*+/+ in *Tg(CMV:GFP-map1lc3b)* and *Tg(mpeg1::mcherry-F)*^*umsF001*^ backgrounds (Zhang *et al*., 2020). Embryos from adult fish were handled and treated pre-infection and post infection as described (Masud *et al*., 2019a).

### Bacterial strains and infection experiments

*S*. Typhimurium SL1344 strains were used for this study, constitutively expressing *mCherry* or the *mCherry* marker in combination with a GFP biosensor for ROS (p*katGp-gfpOVA)* (Burton *et al*., 2014). Culturing of bacteria, preparation of infection inocula and infection delivery were performed as described before (Masud *et al*., 2019a). Briefly, bacteria were resuspended in PBS supplemented with 2% polyvinylpyrrolidone-40 (PVP) to obtain the low dose (200-400 CFU, for survival curves and CFU counts experiments) or high dose (2000-4000 CFU, for imaging experiments). Bacterial inoculum was injected systemically into the caudal vein of 2 dpf anaesthetized embryos. Survival of infected larvae was recorded at 24 hour intervals.

### Determination of *in vivo* bacterial (CFU) counts

The *in vivo* bacterial counts were determined as described in (Masud *et al*., 2019a). Briefly, homogenized infected embryos were serially diluted and plated out on solid media to enumerate bacterial colonies.

### Dram1 knockdown and overexpression experiments

Dram1 expression was altered by injecting 1 nl of a 0.1 mM solution of a previously described antisense morpholino MO1-dram1 (AAGGCTGGAAAACAAACGTACAGTA) or by injecting at a concentration of 100 pico grams of *dram1* mRNA in 1 nl (van der Vaart et al., 2014).

### Image acquisition and image analysis

Infected embryos were fixed at 4 hpi for image acquisitions over the yolk sac region. A 63x water immersion objective (NA 1.2) with a Leica TCS SPE system was used. For quantification of GFP-Lc3-*Salmonella* associations and ROS biosensor activation, the images acquired were analyzed through Z-stacks in Leica LAS AF Lite software and bacterial clusters were observed and manually counted in the overlay channel as previously described (Masud *et al*., 2019a). Max projections in the overlay channels were used for representative images.

### Statistical analysis

All data sets were analyzed with Prism 7 software. Survival curves were analyzed with Log rank (Mantel-Cox) test. For CFU counts, one-way ANOVA was performed on Log-transformed data. Data of GFP-Lc3-positive infected phagocytes and biosensor-positive phagocytes were analyzed with unpaired parametric t-test between two groups and for multiple groups the one-way ANOVA test was performed and corrected for multiple comparisons.

## Acknowledgements

We thank Dirk Bumann (University of Basel) for sharing of the *Salmonella* strains used in this study, and Daniel Klionsky (University of Michigan) for the zebrafish transgenic line. We are also grateful to all members of the fish facility team for zebrafish care. S.M. was supported by a fellowship from the Higher Education Commission of Pakistan and the Bahaudin Zakriya University, Multan. R.Z. was supported by a fellowship from the China Scholarship Council and T.K.P. by an individual Marie Curie fellowship (PIEF-GA-2013-625975).

